# Impact of cryopreservation on immune cell metabolism as measured by SCENITH

**DOI:** 10.1101/2024.06.12.598758

**Authors:** Curtis Luscombe, Eben Jones, Michaela Gregorova, Nicholas Jones, Laura Rivino

**Author notes:** Corresponding author Mailing address: School of Cellular and Molecular Medicine, University of Bristol, Bristol, United Kingdom. These authors contributed equally to this work (Curtis Luscombe, Eben Jones).

## Abstract

The dynamic functioning of immune cells is regulated by cellular metabolic processes, and there is growing interest in the study of immunometabolic correlates of dysfunctional immune responses. SCENITH is a novel flow cytometry-based technique that allows for *ex vivo* metabolic profiling of immune cells within heterogeneous samples. Cryopreservation of clinical samples is frequently undertaken to facilitate high throughput processing, but is thought to lead to cellular metabolic dysfunction. We aimed to investigate the impact of cryopreservation on immune cell metabolism, harnessing SCENITH’s unique ability to describe the divergent bioenergetic characteristics of distinct immune cell subsets.

We demonstrate that T cells undergoing activation with a CD3/CD28 stimulus are less readily metabolically reprogrammed following cryopreservation. Additionally, we find that cryopreservation introduces a time-dependent metabolic artefact that favours glycolysis and impairs oxidative phosphorylation, suggesting that cryopreservation results in mitochondrial dysfunction. Despite this artefact, SCENITH was still able to reveal the distinct bioenergetic profiles of contrasting immune cells populations following cryopreservation – for example, non-classical monocytes have a higher mitochondrial dependence than classical monocytes, and T cell CD69 expression is associated with an upregulation of glycolytic capacity.

Whilst we believe that SCENITH can provide valuable information about immune cell metabolism even in cryopreserved samples, our findings have important implications for the design of future studies. Investigators should carefully consider how to process and store clinical samples to ensure that cryopreservation does not confound analyses, particularly where longitudinal sampling is required.

## Introduction

Immune cell behaviour is underpinned by tightly regulated cellular metabolic processes, and metabolic pathway intermediaries are directly implicated in epigenetic changes that are necessary for the dynamic functioning of immune cells[1]. Naïve T cells maintain low levels of transcription and translation, largely meeting their limited bioenergetic requirements by catabolising glucose and amino acids via oxidative phosphorylation[2]. Conversely, T cell activation, differentiation, and effector functions are associated with higher overall cellular energetic expenditure, which is achieved by broadly upregulating cellular metabolism, and aerobic glycolysis in particular[3-5]. A deeper understanding of the immunometabolic correlates of physiological and dysfunctional adaptive immune responses in health and disease offers the potential to identify novel immunomodulatory therapeutic targets in the treatment of infection, auto-immune disease, and cancer.

Single Cell ENergetIc metabolism by profiling Translation inHibition (SCENITH) is a novel functional assay which can be used to metabolically profile individual immune cells using flow cytometry. Compared to bulk analysis techniques such as extracellular flux analysis (Seahorse), SCENITH can facilitate a comparison between distinct cell populations in mixed samples (e.g. whole blood), including those that occur at a relatively low frequency. Furthermore, SCENITH does not require purification of immune cells for metabolic profiling, which is highly advantageous when analysing clinical samples which may be of a limited volume. Protein translation is an energetically expensive process, the rate of which is closely correlated with the overall rate of adenosine triphosphate (ATP) synthesis in immune cells[6]. In SCENITH, cells are treated with puromycin, which inhibits protein synthesis by becoming incorporated into – and thus causing the premature termination of – nascent polypeptide chains[7]. The rate of puromycin incorporation is quantified by staining cells with anti-puromycin antibody before measuring its mean fluorescence intensity (MFI). Anti-puromycin MFI acts as the assay’s readout and acts as a proxy measure for the cell’s overall energetic expenditure[6].

Additionally, SCENITH takes advantage of selective inhibition of specific metabolic pathways that contribute to cellular biosynthesis of ATP (Fig. 1). 2-deoxy-D-glucose (2DG) is a glucose analogue that inhibits hexokinase (HK) and thus glycolysis[8]. Oligomycin-A (OMA) is an ATP synthase (ATPS) inhibitor which inhibits oxidative phosphorylation[9]. Comparing the anti-puromycin MFI of cells treated in parallel with 2DG, OMA, or both (‘DGO’) versus an uninhibited positive control (‘P-Co’) facilitates the calculation of a cell’s reliance upon glycolysis, oxidative phosphorylation, or other fuels (the oxidation of fatty acids and amino acids, Fig. 2).

**Figure 1.**
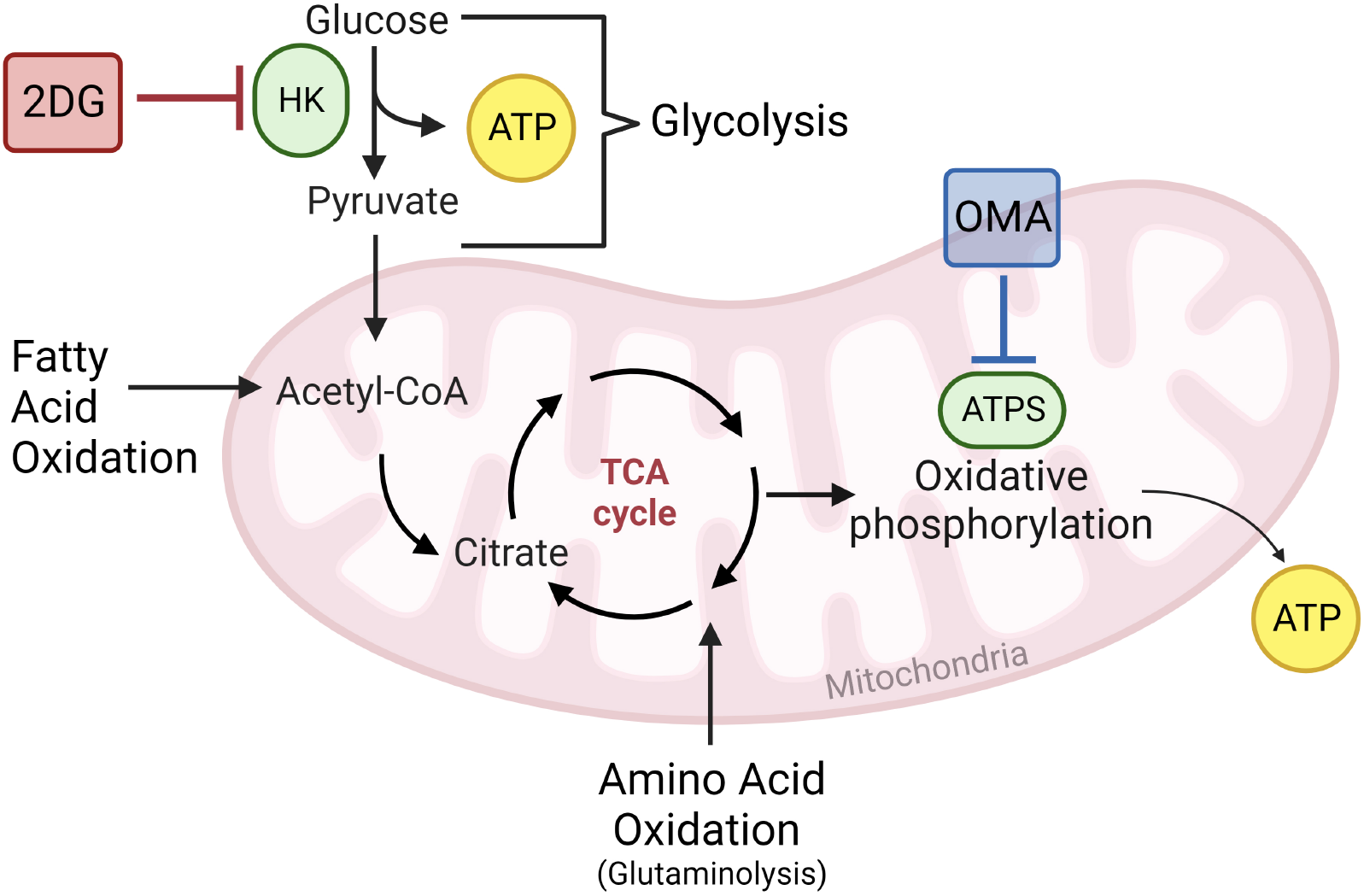
Schematic representation of the cellular bioenergetic pathways that contribute to ATP biosynthesis. In glycolysis, a sequence of reactions sees glucose converted into pyruvate, resulting in the generation of ATP (yellow). Pyruvate passes into the mitochondria, where it is converted into Acetyl-CoA before feeding into the tricarboxylic acid (TCA) cycle. A further sequence of reactions within this cycle leads to the activation of electron carrier molecules, which deliver electrons to the electron transport chain, resulting in ATP generation via oxidative phosphorylation. Fatty acids and amino acids are oxidised and also feed into the TCA cycle, thus also generating ATP. 2-deoxy-D-glucose (2DG, red) inhibits hexokinase (HK, green), therefore inhibiting glycolysis. Oligomycin A (OMA, blue) inhibits ATP synthase (ATPS, green), therefore inhibiting oxidative phosphorylation. Produced using BioRender.

**Figure 2.**
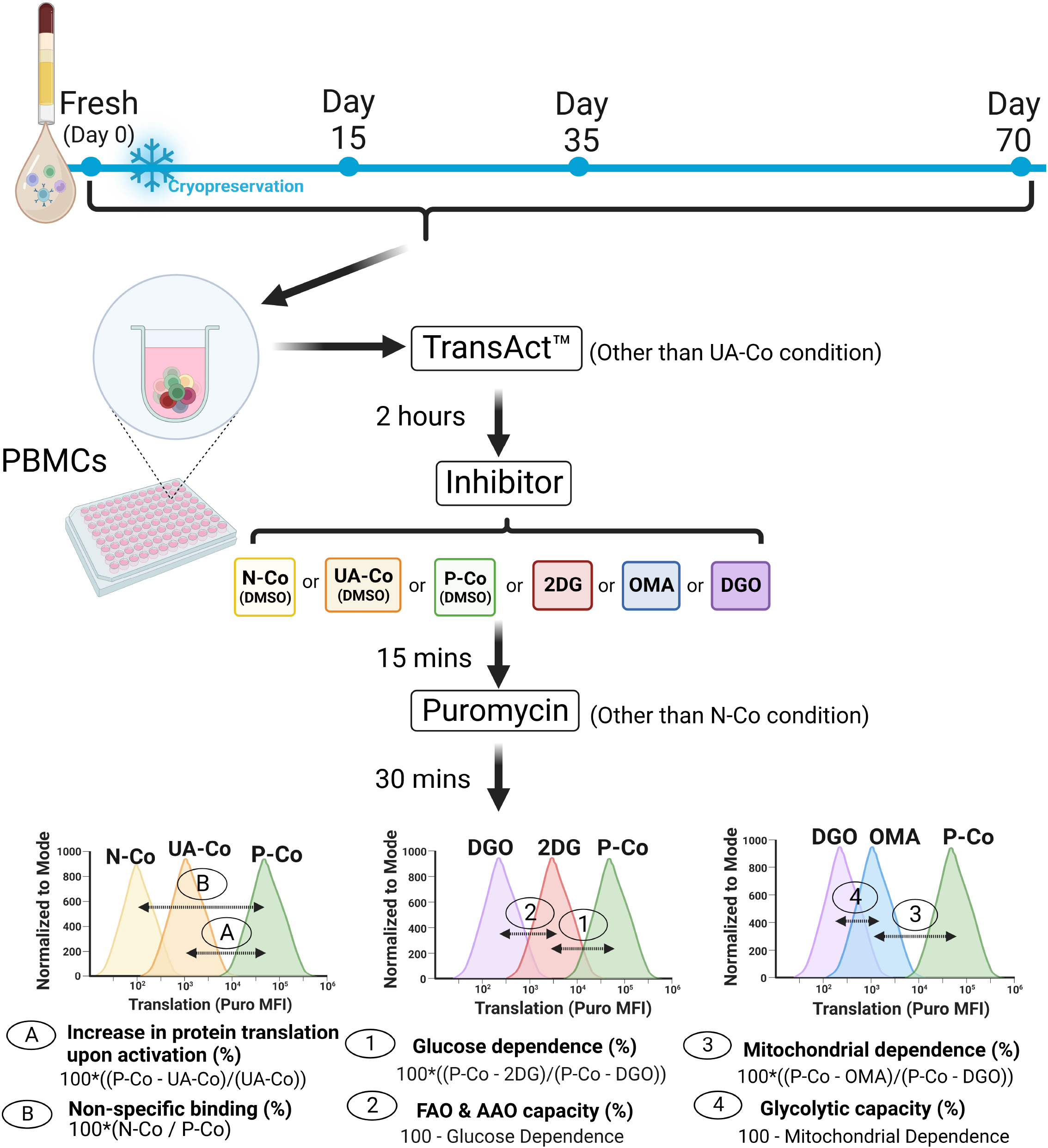
Schematic of cryopreservation study design and SCENITH calculations. PBMCs were taken from selected timepoint and incubated first with TransAct anti-CD3/CD28 agonist, then with metabolic inhibitors (or DMSO), then with puromycin. Controls and treatment conditions are defined in Table 1. Produced using BioRender.

Argüello *et al*. originally describe performing SCENITH on whole blood, and it has subsequently been applied to peripheral blood mononuclear cell (PBMC) samples[10, 11], including samples which have been cryopreserved[12]. Cryopreservation of PBMCs prior to their use in phenotypic and functional assays allows for the biobanking of samples, and for cheaper, more practical, and higher-throughput processing of multiple samples in parallel. However, environmental changes directly influence cellular bioenergetics[13], meaning that cryopreservation may introduce a systematic bias that compromises the reliability and reproducibility of metabolic assays.

**Table 1.** Definition of experimental conditions used in this study:

| Condition: | Abbreviation: | Anti-CD3/CD28 agonist: | Metabolic inhibitor: | Puromycin: |
| --- | --- | --- | --- | --- |
| No puromycin negative control | N-Co | Y | DMSO | N |
| Unactivated control | UA-Co | N | DMSO | Y |
| Positive control | P-Co | Y | DMSO | Y |
| 2DG | 2DG | Y | 2DG | Y |
| OMA | OMA | Y | OMA | Y |
| DGO | DGO | Y | 2DG & OMA | Y |

The impact of cryopreservation on cellular bioenergetics measured via the Seahorse assay has previously revealed progressive mitochondrial dysfunction and a shift towards dependence upon glycolysis[14, 15]. However, these studies analysed mixed PBMC samples using a bulk analysis assay. Given that distinct immune cell populations have divergent bioenergetic characteristics, the conclusions that can be drawn from the existing literature are therefore inherently limited. In this study, we aim to investigate the impact of cryopreservation on immune cell metabolism as measured by SCENITH, with a particular focus on T cells.

## Results

### The effect of cryopreservation on immune cell recovery, immune cell viability and sample composition

PBMCs were analysed immediately after isolation (fresh) and after 15, 35 and 70 days of cryopreservation. Cellular recovery and viability were calculated at each timepoint immediately after thawing using the following equations:

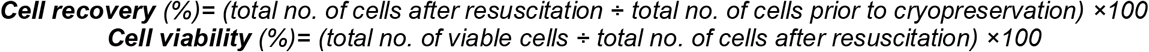

As expected, the cellular recovery (Fig. 3A) and viability (Fig. 3B) of PBMCs decreased after cryopreservation and thawing. After 35 days, there was a significant decrease in the percentage recovery (p=0.0197) and viability (p=0.0270) when compared with fresh PBMCs, a trend which continued at 70 days. The lowest mean recovery was 60.24±0.89% on day 70 (Fig. 3A) whilst the lowest mean viability was 82.90±1.60% on day 35 (Fig. 3B).

**Figure 3.**
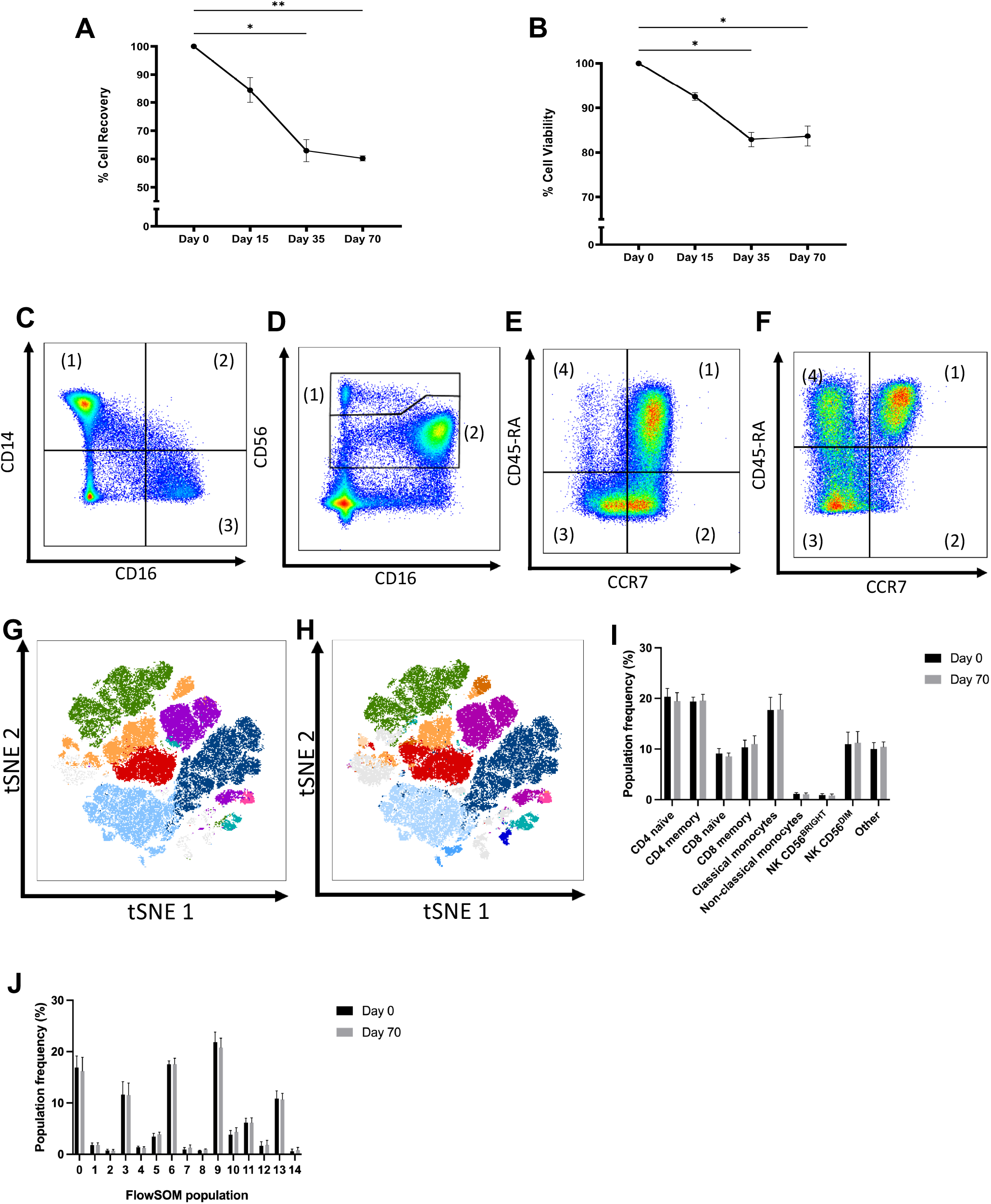
Cell recovery, viability and composition before and after cryopreservation. Human PBMCs were cryopreserved and thawed at selected time points. Cell recovery (**A**) and viability (**B**) were calculated for each time point (*\*P<0*.*05, **P<0*.*01, Dunn’s multiple comparisons test comparing Days 15, 35 and 70 to fresh only*). CD3-, CD56- cells were gated according to their expression of CD16 and CD14, establishing classical (1), intermediate (2) and non-classical (3) monocyte subsets **(C)**. CD3-, CD56+ cells were gated according to their expression of CD16 and CD56, establishing NK CD56^Bright^ (1) and NK CD56^Dim^ (2) NK cell subsets **(D)**. CD3^+^, CD4^+^ cells were gated according to their expression of CCR7 and CD45-RA, establishing naïve (1), TCM (2), TEM (3), and TEMRA CD4^+^ T cell subsets **(E)**. CD3^+^, CD8^+^ cells were gated according to their expression of CCR7 and CD45- RA, establishing naïve (1), TCM (2), TEM (3), and TEMRA CD8^+^ T cell subsets **(F)**. t-SNE demonstrating cellular composition of resting PBMCs at day 0 were superimposed with manually gated populations **(G)**, or clusters identified using the FlowSOM algorithm **(H)**, or with manually gated populations (**H**). Frequency of each manually gated (**I**) or FlowSOM (**J**) population at time point 0 and 70 (*Wilcoxon matched-pairs signed rank test comparing fresh to Day 70, n=5)*.

The impact of cryopreservation on PBMC cellular composition was assessed by analysing cell viability on manually-gated subpopulations, specifically: classical, intermediate and non- classical monocytes (respectively CD14^+^ CD16^-^, CD14^+^ CD16^+^ and CD14^-^ CD16^+^; Fig. 3C), CD56^Bright^ and CD56^Dim^ NK cells (Fig. 3D), CD4^+^ T cell subsets (Fig. 3E) and CD8^+^ T cell subsets (Fig. 3F). A full gating strategy is outlined in Supplementary Fig. 1. Additionally, the FlowSOM clustering algorithm was used to identify 15 distinct populations within PBMC samples based on cellular expression of lineage and T cell differentiation markers (CCR7, CD45RA) within our panel (Supplementary Fig. 2). Manually gated (Fig. 3G) and FlowSOM (Fig. 3H) populations were visualised by superimposing them onto a map created using the unsupervised nonlinear dimensionality reduction algorithm t-Distributed Stochastic Neighbour Embedding (tSNE)[17]. There is a strong visual correspondence between FlowSOM clusters and manually gated populations, and between tSNE and FlowSOM clustering (Fig. 3G-H, Table 2). There was no significant difference in the frequency in either manually gated (Fig. 3I) or FlowSOM populations (Fig. 3J) at fresh and day 70, indicating that cryopreservation universally decreased PBMC viability but did not impact upon PBMC composition in this study.

**Table 2.** Frequencies and descriptions of manually gated and FlowSOM populations:

| Fig. 2B colour: | Manually gated population: | Manually gated population frequency (%): | FlowSOM population: | FlowSOM population frequency (%): | Fig. 2B colour: |
| --- | --- | --- | --- | --- | --- |
|  | Classical monocyte | 17.8 | 0 | 14.6 |  |
|  | Non-classical monocyte | 1.10 | 1 | 1.83 |  |
|  | NK CD56<sup>bright</sup> | 9.87 | 2 | 9.78 |  |
|  | NK CD56<sup>dim</sup> | 11.1 | 4 | 11.6 |  |
|  | CD4<sup>+</sup> naïve | 18.6 | 6 | 21.3 |  |
|  | CD4<sup>+</sup> Memory | 18.5 | 5 | 17.5 |  |
|  |  |  | 7 | 1.12 |  |
|  |  |  | 8 | 0.85 |  |
|  | CD8<sup>+</sup> naïve | 1.83 | 13 | 16.7 |  |
|  | CD8<sup>+</sup> Memory | 10.7 | 11 | 6.17 |  |
|  |  |  | 12 | 1.77 |  |
|  |  |  | 14 | 9.71 |  |

### The effect of cryopreservation on the T-cell translational response to activation

PBMCs were incubated for 2 hours with (P-Co/N-Co) or without (UA-Co) the addition of CD3/CD28 agonist TransAct, and then treated with (P-Co/UA-Co) or without (N-Co) puromycin. The background non-specific binding was calculated using the following equation:

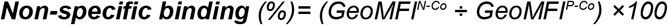

A general trend of increased non-specific binding was demonstrated at day 70 when compared to fresh (Fig. 4A), but this was only statistically significant within the CD4^+^ naïve (p=0.0099) and classical monocyte (p=0.0099) populations (Fig. 4A). The non-specific binding was significantly different between NK CD56^Dim^ and CD8^+^ Naïve (p=0.0021), NK CD56^Dim^ and classical monocytes (p=0.0003) and CD8^+^ TEM and classical monocytes (p=0.0161) at fresh, however these significant differences were not found at day 70 (Fig. 4A). Together, these results indicate that the degree of non-specific binding varies according to cell type and may be influenced by factors such as cryopreservation, highlighting the value of including a no-puromycin negative control (N-Co) when performing SCENITH.

**Figure 4.**
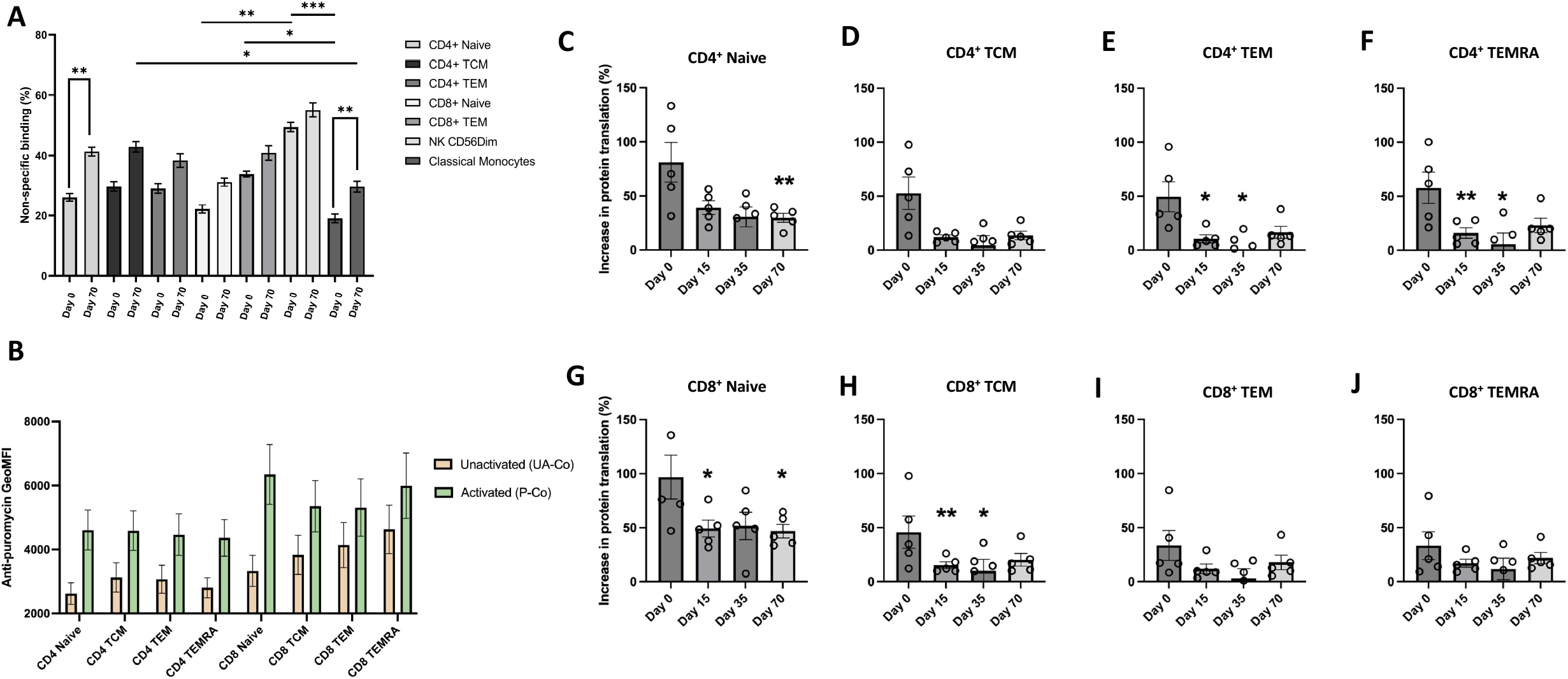
Response of T cell subsets before and after cryopreservation. Human PBMCs were incubated with CD3/CD28 agonist TransAct. Non-specific binding was compared between the seven most abundant populations at fresh and day 70 (**A**). Anti- puromycin GeoMFI for T cell subsets in unactivated and activated conditions was compared at fresh (**B**). Comparing unactivated (UA-Co) and activated (P-Co) samples, increase in protein translation upon activation is shown for CD4^+^ Naïve **(C)**, TCM (**D**), TEM (**E**), TEMRA (**F**), and CD8^+^ Naïve (**G**), TCM (**H**), TEM (**I**), TEMRA (**J**) T-cells. (*\*P<0*.*05, **P<0*.*01, Friedman Test with Dunn’s multiple comparisons test comparing Days 15, 35 and 70 to fresh only, n=5*).

At fresh, resting and activated CD8^+^ T cell subsets were uniformly more metabolically active than CD4^+^ T cell subsets. In resting CD4^+^ and CD8^+^ cells (UA-Co), naïve subsets were less metabolically active than memory subsets. After activation (P-Co), the metabolic activity of all subsets increased, with CD4^+^ and CD8^+^ naïve subsets more metabolically active than their respective memory subsets, indicating that despite their lower baseline activity, the metabolic activity of naïve cells is more rapidly upregulated than memory cells following 2 hours of activation with an anti-CD3/CD28 agonist (Fig. 4B).

The increase in protein translation in response to activation was calculated by comparing protein translation in unactivated (UA-Co) and activated (P-Co) conditions at each timepoint, using the following equation:

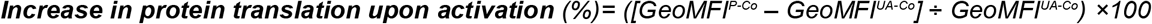

There was a general decrease in the percentage translational response after cryopreservation for all T cell subpopulations (Fig. 4C-4J). Compared to fresh, there was a significant decrease in the percentage increase of translation after activation within the CD4^+^ Naïve T-cells at day 70 (p=0.0099) (Fig. 4C), CD4^+^ TEM at day 15 (p=0.0212) and day 35 (p=0.0212) (Fig. 4E), CD4^+^ TEMRA cells at day 15 (p=0.0099) and day 35 (p=0.0429) (Fig. 4F), CD8^+^ Naïve T-cells at day 15 (p=0.0429) and day 70 (p=0.0212) (Fig. 4G) and CD8^+^ TCM cells at day 15 (p=0.0044) and day 35 (p=0.0429) (Fig. 4H).

After 2 hours of activation, CD4^+^ Naïve cells increased translation by 81.1% at fresh (Fig. 4C), whilst CD4^+^ memory cells increase translation by approximately 50% (Fig. 4D-4F). This decreased to 29.9% for CD4^+^ naïve and approximately 15% for the CD4^+^ memory populations at day 70, equivalent to 36% and 30% of the fresh values, respectively.

After 2 hours of activation, CD8^+^ naïve cells increased translation by 96.9% at fresh (Fig. 4G), whilst CD8^+^ memory cells increase by approximately 37% (Fig. 4H-J). This decreases to 46.9% and approximately 20% at day 70, equivalent to 48.4% and 54% of the fresh values, respectively.

Within both CD4^+^ and CD8^+^ subsets, naïve populations showed the greatest increase in translation (Fig. 4C&4G), whilst TEM populations show the lowest increase in translation (Fig. 4E&4I), both at fresh and at day 70. However, there is a greater decrease in the translational response to activation following cryopreservation found for CD4^+^ subpopulations than for CD8^+^ subpopulations.

### Metabolic characteristics of immune cell subsets, and the effect of cryopreservation on immune cell inhibition by 2-Deoxy-D-Glucose and Oligomycin A

The metabolic characteristics of the highest frequency immune cell subsets were compared at fresh using the following equations, taking into account non-specific binding (N-Co, see Fig. 4):

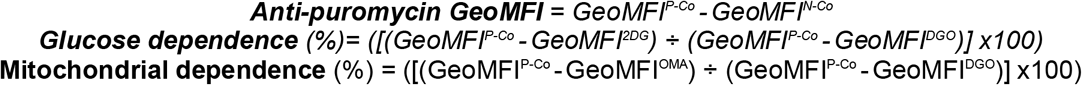

At fresh, classical monocytes were the most metabolically active immune cell subset, undertaking approximately twice as much protein translation as non-classical monocytes (mean anti-puromycin GeoMFI of 14075 vs 6969, respectively), with lymphocyte subsets less active still, potentially reflecting their smaller size in comparison to monocytes (mean anti-puromycin GeoMFI ranging from 3262 (CD4^+^ TCM) to 4987 (CD8^+^ naïve), Fig. 5A).

**Figure 5.**
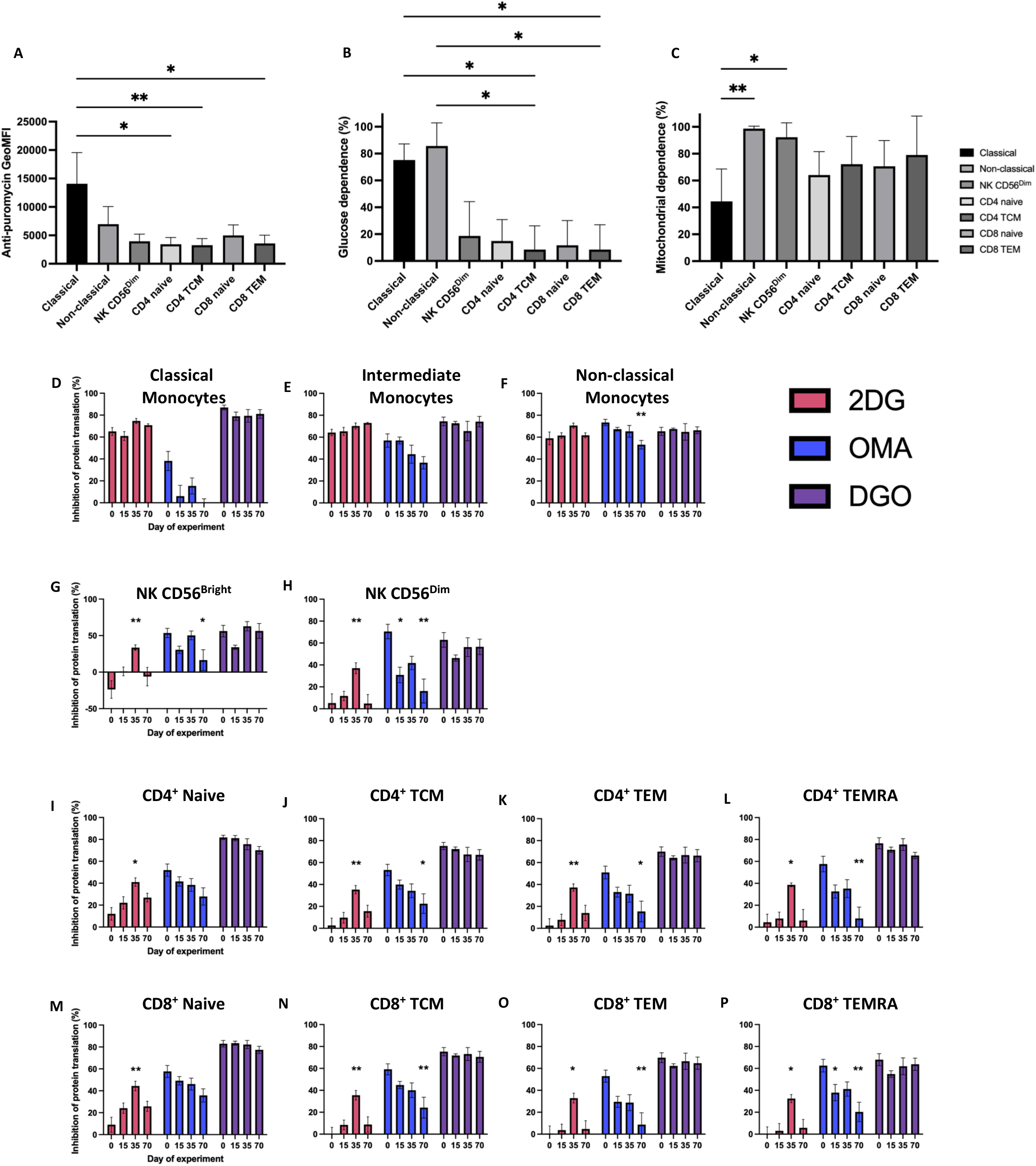
Metabolic characteristics of immune cell subsets at fresh, and inhibition of protein translation following treatment with 2DG, OMA, or DGO at different timepoints. For manually gated subsets, anti-puromycin GeoMFI at fresh is shown to indicate overall levels of metabolic activity (**A**). Glucose dependence (**B**) and mitochondrial dependence (**C**) are also shown. (*\*P<0*.*05, **P<0*.*01, Kruskall-Wallis Test with Dunn’s multiple comparisons test comparing specified subsets, n=5)*. Next, anti-puromycin GeoMFI was calculated for PBMCs treated with 2DG (red), OMA (blue), or DGO (purple), and then compared to the P-Co anti-puromycin GeoMFI. The inhibition of protein translation exerted by each treatment condition was then calculated as a percentage, with non-specific binding taken into account. Inhibition of in protein translation shown for Classical (**D**), Intermediate (**E**) & Non-Classical Monocytes (**F**), NK CD56^Bright^ (**G**) & NK CD56^Dim^ (**H**) cells, CD4^+^ Naïve (**I**), TCM (**J**), TEM (**K**), TEMRA (**L**), and CD8^+^ Naïve (**M**), TCM (**N**), TEM (**O**), TEMRA (**P**) T-cells. (*\*P<0*.*05, **P<0*.*01, Friedman Test with Dunn’s multiple comparisons test comparing Days 15, 35 and 70 to fresh only, n=5*).

Additionally, both classical and non-classical monocyte populations had the highest mean levels of glucose dependence (75 and 86%, respectively), whereas lymphocyte subsets had mean glucose dependencies of under 20% (Fig. 5B). When considering mitochondrial dependence, further differences between classical and non-classical monocytes became apparent, with classical monocytes having a significantly lower mean mitochondrial dependence (45 vs 99% respectively, p=0.0091, Fig. 5C). Compared to other lymphocyte subsets, NK CD56^Dim^ had a higher mitochondrial dependence (<75 vs 92%), potentially reflecting the activation and subsequent switch to aerobic glycolysis for non-NK cell lymphocyte subsets following the addition of TransAct. When considered together, these three different indicators of immune cell metabolic activity can clearly delineate between the manually gated subsets specified by our panel, with other T cell subsets showing comparable patterns (data not shown).

Next, the impact of cryopreservation upon the inhibitory effect of 2DG, OMA or both (DGO) was considered for each manually gated subset, using the following equation:

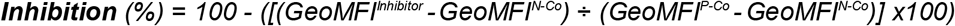

Firstly, there was no statistically significant difference in the inhibitory activity of the DGO combination at any timepoint and for any cell subset (Fig. 5D-5P), indicating that the basal level of inhibition was unchanged by cryopreservation.

OMA demonstrated a trend towards decreased inhibitory activity in all cell subsets as the duration of cryopreservation increased (Fig. 5D-5P). Compared to fresh, the inhibitory activity of OMA was significantly reduced at day 70 in non-classical monocytes (p=0.0018) (Fig. 5F), NK CD56^Bright^ (p=0.0212) (Fig. 5G), NK CD56^Dim^ (p=0.0018) (Fig. 5H), CD4^+^ TCM (p=0.0429) (Fig. 5J), CD4^+^ TEM (p=0.0212) (Fig. 5K), CD4^+^ TEMRA (p=0.0044) (Fig. 5L), CD8^+^ TCM (p=0.0044) (Fig. 5N), CD8^+^ TEM (p=0.0044) (Fig. 5O) and CD8^+^ TEMRA (p=0.0018) (Fig. 5P) cells.

2DG demonstrated a trend towards increased inhibitory activity in the lymphocyte populations (Fig. 5G-5P), while maintaining a consistent percentage inhibition in the monocyte populations (Fig. 5D-5F). In lymphocytes, 2DG inhibition was maximal at day 35 (Fig. 5I-5P). Compared to fresh, the inhibitory activity of 2DG was significantly increased at day 35 in NK CD56^Bright^ (p=0.0099) (Fig. 5G), NK CD56^Dim^ (p=0.0099) (Fig. 5H), CD4^+^ Naïve (p=0.0212) (Fig. 5I), CD4^+^ TCM (p=0.0099) (Fig. 5J), CD4^+^ TEM (p=0.0099) (Fig. 5K), CD4^+^ TEMRA (p=0.0429) (Fig. 5L), CD8^+^ Naïve (p=0.0099) (Fig. 5M), CD8^+^ TCM (p=0.0099) (Fig. 5N), CD8^+^ TEM (p=0.0429) (Fig. 5O) and CD8^+^ TEMRA (p=0.0212) (Fig. 5P) cells.

Despite the artefact introduced by cryopreservation, the metabolic profiles of classical vs non-classical monocytes remained distinct following cryopreservation and followed the same pattern as when analysed fresh, with classical monocytes having higher levels of protein translation (anti-puromycin GeoMFI 14897 vs 10569, p=0.055556), similar levels of glucose dependence (87 vs 92%, p=0.746032), and lower levels of mitochondrial dependence (7 vs 81%, p= 0.007937, data not shown).

### The metabolic differences observed between CD69+ and CD69- T cells are preserved following cryopreservation

As a cytometry-based technique, a core strength of SCENITH compared to bulk analysis assays is the ability to compare between cells subsets within heterogeneous samples (e.g. PBMCs) based upon cellular expression of phenotypic markers. CD69 is an early activation marker expressed by T cells and is associated with divergent immune cell metabolic signatures when measured via other single-cell techniques[18]. We therefore asked whether CD69 expression was associated with divergent T cell metabolic profiles when measured via SCENITH, and whether this was still evident following cryopreservation.

PBMCs were incubated for 2 hours with the addition of CD3/CD28 agonist TransAct, and then treated with metabolic inhibitors as described previously (Fig. 2). CD69^+^ and CD69^-^ populations were manually gated on (Fig. 6A). Subsequently, the glycolytic capacity and glucose dependencies of CD69^+^ and CD69^−^ populations were established, and the difference between CD69^+^ and CD69^-^ populations was defined using the following equations:

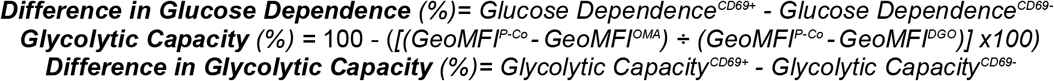

**Figure 6.**
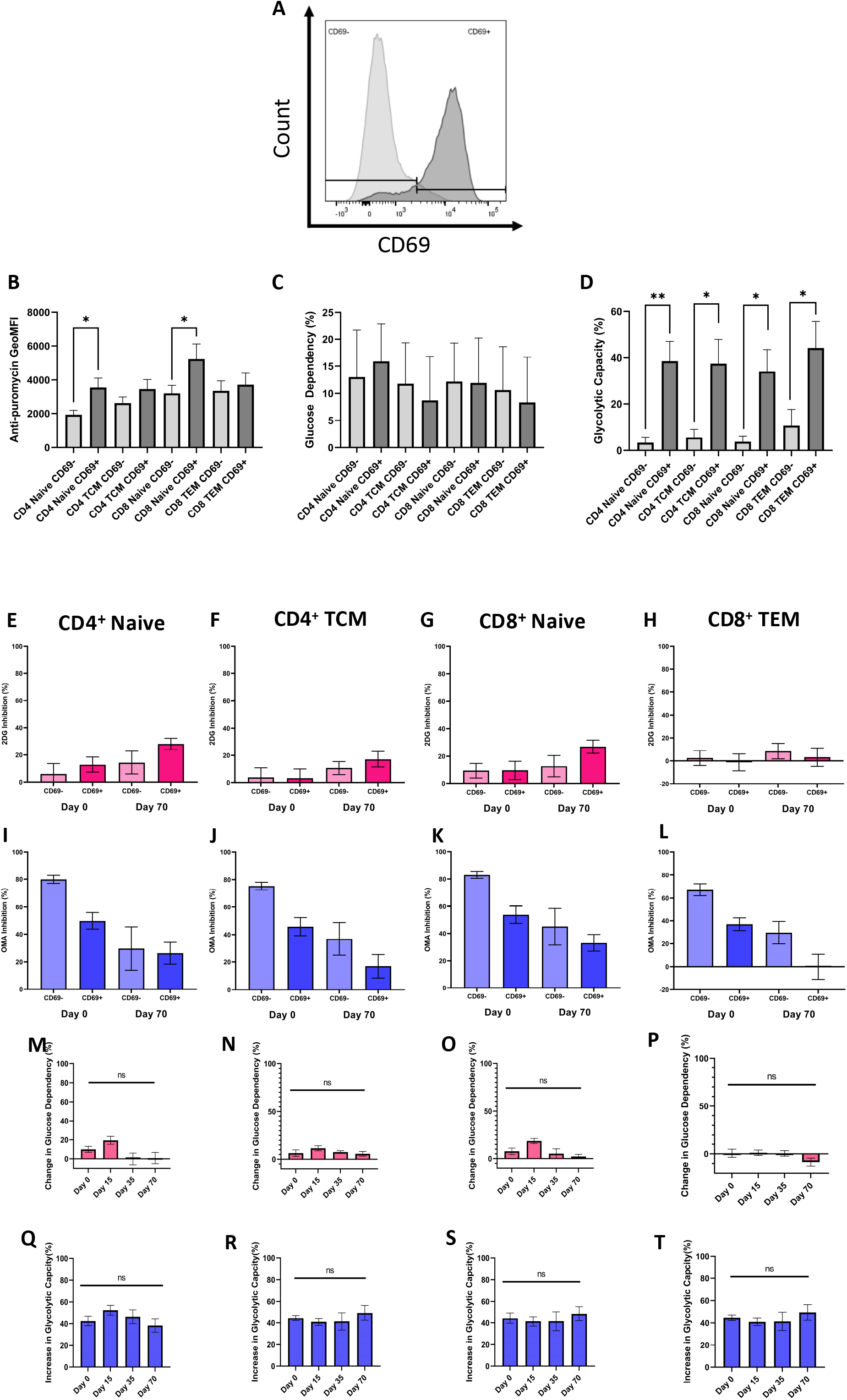
Metabolic differences observed between CD69^+^ and CD69- T cells are preserved following cryopreservation. CD3^+^ T cell subpopulations were gated according to their expression of CD69 (dark grey), as determined by the unactivated control (light grey) (**A**). Baseline (fresh), for CD69+ and CD69- subsets, anti-puromycin GeoMFI at fresh is shown to indicate overall levels of metabolic activity (B). Glucose dependence (C) and glycolytic capacity (D) are also shown. (*P<0.05, **P<0.01, Mann-Whitney U tests comparing specified CD69+ and CD69- populations withinin subsets, n=5). 2DG inhibition was determined for the CD4^+^ Naïve (E), CD4^+^ TCM (F), CD8^+^ Naïve (G), CD8^+^ TEM (H) and OMA inhibition for the CD4^+^ Naïve (I), CD4^+^ TCM (J), CD8^+^ Naïve (K), CD8^+^ TEM (L) CD69- and CD69+ subpopulations at day 0 (fresh) and day 70. Following this, the difference in glucose dependency for the CD4^+^ Naïve (M), CD4^+^ TCM (N), CD8^+^ Naïve (O), CD8^+^ TEM (P) and difference in glycolytic capacity for the CD4^+^ Naïve (Q), CD4^+^ TCM (R), CD8^+^ Naïve (S), CD8^+^ TEM (T) CD69- and CD69+ subpopulations at day 0 (fresh) and day 70. (ns*P>0*.*05, Dunn’s multiple comparisons test comparing Days 15, 35 and 70 to fresh only, n=5*).

CD69^+^ and CD69^-^ populations of the naïve and most abundant memory subset of CD4^+^ and CD8^+^ T cells are compared at fresh. Within the naïve CD4^+^ and CD8^+^ populations, CD69 expression was associated with a significant increase in protein translation, indicating increased metabolic activity (anti-puromycin GeoMFI of 1932 vs 3550 and 3205 vs 5234, p=0.0159 and 0.0317 respectively, Fig. 6B). Within memory CD4^+^ and CD8^+^ populations, CD69 expression was also associated with a trend towards increased protein translation, but this difference was not statistically significant (2620 vs 3459 and 3353 vs 3718, p=0.3095 and 0.8413 respectively, Fig. 6B). CD69 expression was not associated with a particular pattern of glucose dependence (Fig. 6C). Conversely, there is a clear trend towards increased glycolytic capacity associated with T cell CD69 expression, including in CD4^+^ naïve (3.38 vs 38.55%, p=0.0079), CD4^+^ TCM (5.65 vs 37.50%, p=0.0317), CD8^+^ naïve (3.80 vs 34.05%, p=0.0159) and CD8^+^ TEM (10.83 vs 44.11%, p=0.0317) populations (Fig. 6D). Together, these results demonstrate that lymphocyte CD69 expression is associated with increased cellular metabolic activity, met by an upregulation of glycolytic capacity. Although CD69 expression was not associated with higher glucose dependence in this study, a longer duration of activation with an anti-CD3/CD28 agonist may produce different results.

Next, we considered the impact of cryopreservation on the metabolic characteristics of CD69+ and CD69- T cell subpopulations, and we observe patterns consistent with those seen for T cell subsets as a whole (Fig. 5I-5P). Following cryopreservation, CD69^+^ and CD69^-^ subpopulations also show a trend towards increased 2DG inhibition (Fig. 6E-6H) and decreased OMA inhibition (Fig. 6I-6L). However, there was no significant change found in the difference in glucose dependency (Fig. 6M-6P) and glycolytic capacity (Fig. 6Q-6T) between CD69^-^ and CD69^+^ cells in any CD4^+^ and CD8^+^ subsets across the timepoints investigated in this study, demonstrating that the metabolic differences observed between CD69^+^ and CD69^-^ T cells are preserved following cryopreservation.

## Discussion

SCENITH is a novel functional assay which can be used to metabolically profile immune cells using flow cytometry. In this study, we demonstrate that although naïve T cells are less metabolically active than memory T cells at rest, they are more readily activated following the addition of an anti-CD3/28 agonist (Fig. 4). Secondly, we demonstrate that classical and non-classical monocytes have divergent metabolic characteristics, with non-classical monocytes being less metabolically active and have a greater mitochondrial dependence (Fig. 5), consistent with prior transcriptomic comparisons of these subsets[19]. Additionally, we show that within a mixed PBMC sample treated with an anti-CD3/28 agonist, CD69 expression is associated with increased metabolic activity, increased glycolytic capacity and diminished mitochondrial dependence (Fig. 6). This demonstrates the ability of SCENITH to reveal the divergent metabolic characteristics of cell subsets within a mixed sample without the need for purification.

Specifically, this study aimed to investigate the impact of cryopreservation on immune cell metabolism as measured by SCENITH, and we present a number of key findings. Firstly, we show that T cells undergoing activation with a CD3/CD28 stimulus are less readily metabolically reprogrammed following cryopreservation (Fig. 4). Secondly, we find that cryopreservation introduces a metabolic artefact that favours glycolysis and impairs oxidative phosphorylation, as indicated by the increasing inhibitory potential of 2DG and diminishing inhibitory potential of OMA following cryopreservation, with our results suggesting that this artefact becomes more pronounced as the duration of cryopreservation increases (Fig. 5). This was observed in all cell types defined by our multiparametric panel and was not explained by a cryopreservation-induced change in sample cellular composition (Fig. 3). Despite this artefact, SCENITH was still able to reveal the distinct bioenergetic profiles of contrasting immune cells populations following cryopreservation, such as classical and non-classical monocytes (Fig. 5), or T cell subsets stratified by CD69 expression (Fig. 6). As such, we believe that SCENITH can provide valuable insights into how cellular metabolism influences immune cell function even after cryopreservation.

This is the first study to investigate the impact of cryopreservation on immune cell bioenergetics using SCENITH, but our findings are in line with previous studies. Utilising the bulk analysis assay Seahorse, other groups have demonstrated that cryopreservation causes a time-dependent impairment of immune cell mitochondrial function, alongside an enhanced glycolytic response[14, 15]. In Seahorse, spare respiratory capacity is a derived measurement that quantifies the ability of cells to generate additional ATP following an increase in energetic demand[20]. This metric becomes impaired following cryopreservation[15], analogous to our finding of diminished T cell translational responses to CD3/CD28 activation (Fig. 4). Currently, robust evidence as to the mechanistic basis for mitochondrial dysfunction following cryopreservation in immune cells is lacking, but the literature focussing on other cell types (e.g. reproductive cells or placental tissue) highlights the possible role of oxidative stress caused by excessive reactive oxygen species generation, or osmotic stress that compromises the integrity of the mitochondrial membrane[21-23].

Our findings have implications for the design of future studies. Upon study conception, investigators should consider whether the metabolic artefact introduced by cryopreservation will impact upon their ability to satisfactorily address their research question. Performing SCENITH on cryopreserved samples may not be appropriate for studies where participants are sampled longitudinally and biobanking and batch analysis is planned. In these circumstances, cryopreservation is likely to confound results and result in spurious findings. Conversely, intra-donor comparisons between conditions from the same cryopreserved sample is likely to result in valid findings. For example, researchers may wish to investigate the metabolic differences between drug-tested *versus* control cells, which should not be problematic if cells have been processed and stored equally.

There are several important limitations associated with this study. Firstly, we sampled a modest number of healthy donors, limiting the generalisability of our findings to other settings. For example, in the context of acute severe infection, we might expect a greater degree of upregulation of glycolysis in certain T cell subsets than was possible to achieve via our protocol with 2 hours of activation with a CD3/CD28 agonist. Applying this study design to disease states such as acute severe infection may therefore yield different results, with metabolically divergent subsets variably susceptible to artefacts induced by cryopreservation. Additionally, the process of incubating cells with culture medium as required to achieve T cell activation may have introduced a metabolic bias compared to analysis directly after thawing. Furthermore, the addition of an anti-CD3/CD28 agonist will have directly activated T cells, but not NK cells or monocytes. It is plausible that the metabolic characteristics of NK cells and monocytes may have been altered by this step. Finally, we were only able to study four time-points, limiting our ability to predict the impact of cryopreservation beyond 70 days.

## Methods

### Study design and participant characteristics

This study was conducted using peripheral blood mononuclear cells (PBMCs) from five consenting healthy participants (*n=5*). Participants were Caucasian, aged 35.2±8.7, and had a body mass index (BMI) within the healthy range. Two participants were male and three were female. Samples were obtained and used under research ethics approval of the Bristol Biobank (NHS Research Ethics Committee Ref 20/WA/0273, use application U0046).

Timepoints of 15, 35 and 70 days were selected pragmatically to provide an indication of metabolic changes induced by an increasing duration of cryopreservation.

### PBMC isolation and cryopreservation

Venous blood was collected from each participant into EDTA vacutainers (BD). This was diluted 1:1 in PBS and then applied to Ficoll-Paque PLUS (Cytiva) at a 3:1 concentration, and centrifuged at 1256 *g* for 25 minutes, with minimum acceleration and no braking. PBMCs were extracted from the interphase and washed with PBS 1% foetal bovine serum (FBS, Gibco) twice. The total number of cells prior to cryopreservation were counted diluted 1:1 in Trypan Blue dye (Lonza), using a haemocytometer. Cells were suspended into 1mL of freezing media containing 90% FBS and 10% Dimethyl sulfoxide (DMSO) (Sigma-Aldrich) and immediately stored at −80°C for 24 hours in a Mr.Frosty container (Nalgene) containing propan-1-ol. The cryovials were then transferred to liquid nitrogen (−196°C) for storage.

### Thawing of PBMCs and assessment of cell recovery and viability

PBMCs were thawed at the appropriate time point by incubating the cryovial in a water bath for 2 minutes at 37 degrees Celsius. The thawed or fresh (on day 0 before cryopreservation) PBMCs were transferred into a fresh falcon tube and gently resuspended in 14mL of human plasma-like media (HPLM) (Gibco) containing 10% dialysed FBS (dFBS) (Gibco). Cells were counted using a haemocytometer, diluted 1:1 with Trypan Blue dye, to determine total number of cells and total number of viable cells post-thawing, from which the cell recovery and viability was calculated.

### SCENITH assay, flow cytometry staining and data analysis

SCENITH was performed as outlined by Argüello *et al*. [6]. PBMCs were plated at between 0.5-1.0 x10^6^ cells per well into 96-well plates. Six conditions − differing in T-cell activation status, addition of metabolic inhibitors and addition of puromycin − were plated simultaneously (Table 1). PBMCs were stimulated using TransAct (Miltenyi Biotec) at 1:300 titre or left to rest in HPLM for 2 hours. PBMCs were treated with DMSO, 100mM 2-Deoxy-D-Glucose (Merck), 1μM Oligomycin A (Sigma-Aldrich) or a combination of both inhibitors, for 15 minutes at 37oC, 5% CO_2_. 11μg/mL puromycin (Sigma-Aldrich) was added to the relevant wells and incubated for 30 minutes. PBMCs were washed with cold PBS and incubated with viability dye Zombie Aqua (Biolegend) for 10 minutes at room temperature. PBMCs were washed with PBS 1% Bovine Serum Albumin (BSA, Sigma Aldrich) and incubated for 30 minutes at 4oC with antibodies diluted in PBS 1% BSA. PBMCs were fixed and permeabilised with Foxp3 kit (eBioscience), as per manufacturer’s instructions. PBMCs were incubated for 60 minutes at 4°C with antibodies for intracellular markers diluted in Perm-wash (eBioscience), including anti-puromycin-AF488 antibody (see Supplementary Table 1 for a full list of antibodies used). PBMCs were analysed using a LSRFortessa™ X-20 (BD) within 24 hours of processing. Metabolic capacities and dependencies were calculated by comparing the anti-puromycin AF488 GeoMFI of different conditions as outlined in Fig. 2.

### Statistical analysis

Flow cytometry data was analysed using FlowJo v10.8.1 and plug-ins FlowSOM v.3.0.18[16] and ClusterExplorer v.1.7.4. Statistical analyses were performed using GraphPad Prism v.9.5.1. Comparisons between timepoints were made using the Friedman test and Dunn’s multiple comparisons test (fresh vs day 15, day 35 and day 70), or using the Wilcoxon matched-pairs signed rank test and two-stage false discovery rate correction (fresh vs day 70 only). Comparisons between immune cell subsets at the same point were made using the Kruskal Wallis test and Dunn’s multiple comparisons test (for 3 or more subsets) or using the Mann-Whitney test (for 2 subsets, e.g. CD69^+^ vs CD69^-^). In text, averages are expressed as mean +/- standard error of the mean. In figures, error bars indicate the standard error of the mean. For derived calculations (dependencies and capacities), occasionally values of <0% or >100% are obtained, conceptually equivalent to values of 0 and 100% respectively. Where this has occurred, raw data has been adjusted accordingly.

## Author contributions

Curtis Luscombe & Eben Jones were involved in conceptualisation, formal analysis, investigation, methodology, writing − original draft, writing − review & editing. Michaela Gregorova was involved in conceptualisation, formal analysis, methodology, supervision, writing − review & editing. Nicholas Jones was involved in methodology and writing − review & editing. Laura Rivino was involved in conceptualisation, formal analysis, methodology, supervision and writing − review & editing.

## Acknowledgements

The authors wish to acknowledge the assistance of Dr Andrew Herman and Poppy Miller, and the University of Bristol Faculty of Biomedical Sciences Flow Cytometry Facility.

## Data availability

The raw files (.fcs) for all flow cytometry data will be deposited in the FlowRepository upon acceptance of this article.

