## Supplementary Figures and Tables for "Impact of cryopreservation on immune cell metabolism as measured by SCENITH"

### Supplementary Table 1:

| | | Antibody | | | Source | Identifier (Cat. No) |
|---|---|---|---|---|---|---|
| Marker | Staining | Clone | Fluorophore | Concentration | | |
| CD3ε | Extracellular | UCHT1 | AF700 | 1:150 | BD Pharmingen | 557943 |
| CD45RA | Extracellular | HI100 | PE | 1:100 | BD Pharmingen | 555489 |
| CD56 | Extracellular | NCAM16.2 | PE-CF594 | 1:100 | BD Pharmingen | 564849 |
| CD69 | Extracellular | FN50 | BUV395 | 1:20 | BD Pharmingen | 564364 |
| CD4 | Extracellular | RPA-T4 | BV605 | 1:50 | Biolegend | 300555 |
| CD8 | Extracellular | SK1 | APC-Cy7 | 1:20 | Biolegend | 344714 |
| CD14 | Extracellular | M5E2 | PE-Cy7 | 1:25 | Biolegend | 301813 |
| CD16 | Extracellular | 3G8 | BV785 | 1:250 | Biolegend | 302045 |
| CD19 | Extracellular | HIB19 | BUV510 | 1:100 | Biolegend | 302241 |
| CCR7 | Extracellular | G043H7 | BV650 | 1:20 | Biolegend | 353233 |
| Granzyme B | Intracellular | QA16A02 | AF647 | 1:50 | Biolegend | 372219 |
| Perforin | Intracellular | B-D48 | PerCP-Cy5.5 | 1:50 | Biolegend | 353314 |
| CD25 | Extracellular | BC96 | BUV737 | 1:50 | eBioscience | 367-0259-42 |
| Puromycin | Intracellular | 12D10 | AF488 | 1:400 | Merck | MABE343-AF488 |

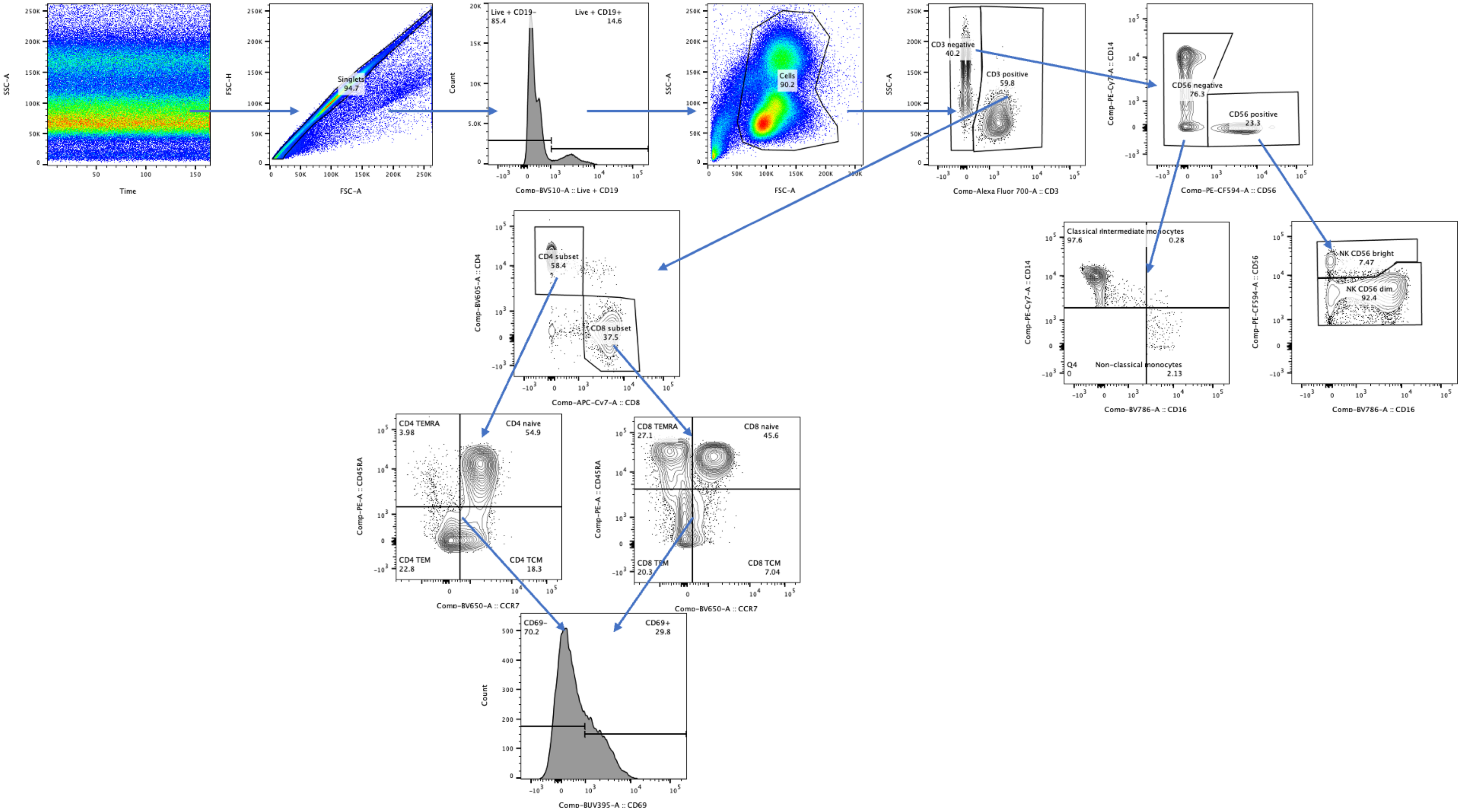

Singlets 94.7
Live + CD19- 85.4
Live + CD19+ 14.6
Cells 90.2
CD3 negative 40.2
CD3 positive 59.8
CD56 negative 76.3
CD56 positive 23.3
Classical monocytes 97.6
Intermediate monocytes 0.28
Q4 0
Non-classical monocytes 2.13
NK CD56 bright 7.47
NK CD56 dim 92.4
CD4 subset 58.4
CD8 subset 37.5
CD4 TEMRA 3.98
CD4 naive 54.9
CD4 TEM 22.8
CD4 TCM 18.3
CD8 TEMRA 27.1
CD8 naive 45.6
CD8 TEM 20.3
CD8 TCM 7.04
CD69- 70.2
CD69+ 29.8
Time
SSC-A
FSC-A
FSC-H
Count
Comp-BV510-A :: Live + CD19
Comp-Alexa Fluor 700-A :: CD3
Comp-PE-Cy7-A :: CD14
Comp-PE-CF594-A :: CD56
Comp-BV786-A :: CD16
Comp-BV605-A :: CD4
Comp-APC-Cy7-A :: CD8
Comp-PE-A :: CD45RA
Comp-BV650-A :: CCR7
Comp-BUV395-A :: CD69

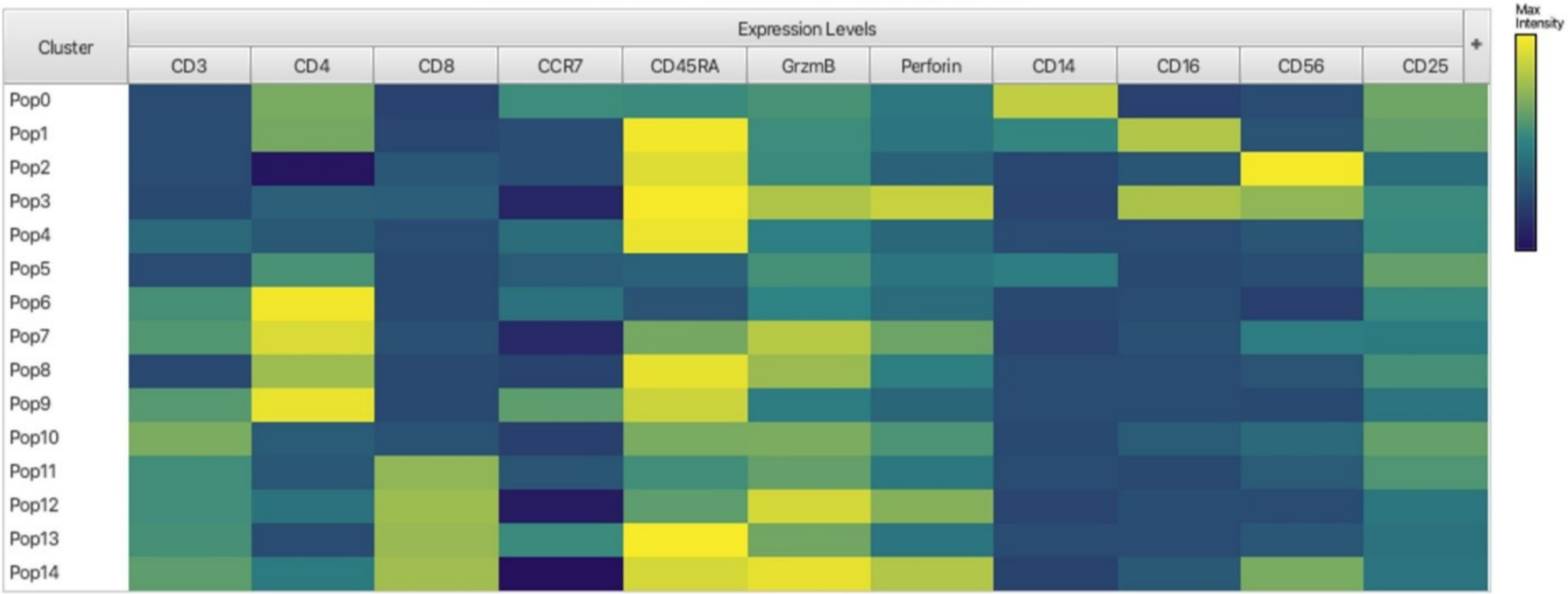

Cluster
Expression Levels
CD3
CD4
CD8
CCR7
CD45RA
GrzmB
Perforin
CD14
CD16
CD56
CD25
Pop0
Pop1
Pop2
Pop3
Pop4
Pop5
Pop6
Pop7
Pop8
Pop9
Pop10
Pop11
Pop12
Pop13
Pop14
Max Intensity
